# Establishment and characterization of LNCaP-TR-BoM, a bone metastasis-derived prostate cancer cell line with an intermediate AR-attenuated phenotype

**DOI:** 10.64898/2026.10.03.756361

**Authors:** Adrian Chao, Xin Lu

## Abstract

**Background:** Progression of prostate cancer under androgen receptor (AR) pathway-targeted therapy spans a phenotypic spectrum, from tumors that sustain AR pathway output to tumors that become independent of AR signaling. Experimental models that occupy the intermediate portion of this spectrum, in which AR is still expressed but pathway output is reduced, are comparatively scarce.

**Methods:** LNCaP cells expressing a triple-modality reporter (LNCaP-TR) were injected into nude mice by the intracardiac route, and a bone lesion that arose in the middle spine of one animal was recovered and propagated as LNCaP-TR-BoM. The subline and its parental line were compared by two-dimensional and three-dimensional culture assays, RNA-seq, qRT-PCR, and immunoblotting. AR pathway output was assessed using *KLK3* expression under AR antagonist treatment. Public prostate cancer datasets provided clinical context. A CRISPR-Cas9 *EPHA3*-edited derivative was evaluated in aggregation and spheroid assays.

**Results:** Relative to parental LNCaP-TR, LNCaP-TR-BoM formed more numerous and larger cell aggregates in monolayer culture and larger, less circular spheroids in both suspension and Matrigel three-dimensional culture. *AR* mRNA was modestly reduced and AR protein remained readily detectable, whereas *KLK3* mRNA and protein were reduced disproportionately more than those of AR. Residual *KLK3* expression remained suppressible by AR inhibitors enzalutamide and darolutamide. Transcriptome-based pseudotime placed LNCaP-TR-BoM between the parental line and the androgen-refractory LNCaP-RF variant. Subsequent differential gene expression (DGE) and pathway enrichment analysis revealed that LNCaP-TR-BoM exhibits transcriptomic rewiring concordant with that of clinical mCRPC, thus evidencing LNCaP-TR-BoM as a potentially clinically relevant model cell line. Finally, *EPHA3* showed the largest fold change among the most significant differentially expressed genes in LNCaP-TR-BoM compared with the parental line. Therefore, *EPHA3* was explored to demonstrate the utility of the LNCaP-TR-BoM cell line as a discovery system for genes that potentially drive the malignancy of bone-metastatic prostate cancer at the intermediate AR-attenuated progression stage. An *EPHA3*-silenced derivative of LNCaP-TR-BoM lacking detectable EPHA3 protein formed smaller aggregates and smaller, more circular spheroids, suggesting *EPHA3* may play a role in driving the malignancy of AR-intermediate bone-metastatic prostate cancer. This finding provides support for the potential utility of the LNCaP-TR-BoM cell line as a discovery system for genes relevant to driving AR-intermediate bone-metastatic prostate cancer.

**Conclusions:** LNCaP-TR-BoM is a bone lesion-derived prostate cancer cell line in which AR expression is retained but AR pathway output is attenuated, accompanied by an mCRPC-concordant transcriptomic shift and altered multicellular organization. It provides a paired discovery system for candidate genes such as *EPHA3*.

## 1. Introduction

Prostate cancer is among the most frequently diagnosed malignancies in men and remains a leading cause of cancer death.^1^ The androgen receptor (AR) signaling axis is the central driver of prostate cancer growth and survival,^2^ and androgen deprivation has formed the backbone of systemic therapy for advanced disease for decades.^3,4^ Potent AR pathway-targeting agents, including the androgen synthesis inhibitor abiraterone and the second-generation AR antagonist enzalutamide, have extended survival in metastatic disease.^5–7^ Resistance nevertheless develops in most patients, and progression is frequently accompanied by metastasis, most often to bone.^8,9^ Bone metastasis and resistance to AR pathway-targeted therapy together account for the majority of prostate cancer mortality.

Resistance is now understood to emerge along a phenotypic spectrum rather than as a single switch.^10,11^ In earlier stages, tumors generally remain AR-positive and sustain pathway output through mechanisms that include *AR* amplification,^12^ ligand-binding domain mutation,^13^ expression of constitutively active splice variants such as AR-V7,^14,15^ intratumoral androgen synthesis,^16^ and sensitization of AR to low ligand concentrations.^17^ Under sustained therapeutic pressure, a clinically important subset of tumors instead undergoes lineage plasticity and transdifferentiates toward phenotypes that have shed AR expression and AR dependence, most prominently neuroendocrine prostate cancer and double-negative prostate cancer.^18–22^ Between these extremes lie tumors in which AR expression or AR transcriptional output is reduced but not extinguished.^11,23,24^ Transcriptomic trajectory analyses of clinical cohorts order samples through such intermediate states before AR-null endpoints,^25^ yet this portion of the spectrum is comparatively understudied, in part because of fewer experimental models.

Most widely used prostate cancer cell lines represent the ends of the spectrum. C4-2 and C4-2B,^26,27^ 22Rv1,^28^ and VCaP^29^ model AR-high disease with sustained pathway output, whereas PC-3,^30^ DU-145,^31^ and LASCPC-01^32^ model AR-null states.^33,34^ Lines that combine derivation from a metastatic site with reduced but persistent AR pathway output are scarce. One established route to more aggressive prostate cancer sublines is *in vivo* selection: recovery of LNCaP-derived tumors from murine hosts yielded the C4-2 and C4-2B lineages with increased tumorigenicity and osseous metastatic propensity,^26,27,35^ and more recent work has defined reproducible preclinical models of AR-driven prostate cancer bone metastasis.^36^

Here we applied *in vivo* selection to generate a model of the intermediate portion of this spectrum. LNCaP cells engineered to express a triple-modality reporter (TR, comprising herpes simplex virus type 1 thymidine kinase, green fluorescent protein, and firefly luciferase^37^) were injected by the intracardiac route, an established experimental-metastasis approach that gives circulating tumor cells access to the skeleton,^38,39^ and lesion development was tracked by bioluminescence imaging. A lesion that arose in the middle spine of one animal was recovered and established in culture as LNCaP-TR-BoM (TR, triple reporter; BoM, bone metastasis-derived). We characterized its growth behavior in two-dimensional and three-dimensional culture, quantified AR expression and AR pathway output, positioned it on a clinical progression trajectory by transcriptome-based pseudotime, compared its transcriptomic rewiring with that of clinical metastatic castration-resistant prostate cancer (mCRPC), and, as a proof of concept for gene discovery, tested the consequences of *EPHA3* disruption.

## 2. Materials and methods

### 2.1. Cell lines and culture

LNCaP, C4-2B and DU-145 were obtained from American Type Culture Collection (CRL-1740, CRL-3315, HTB-81). LNCaP-TR was generated by constitutively expressing a triple-modality reporter fusion protein comprising herpes simplex virus type 1 thymidine kinase, green fluorescent protein, and firefly luciferase.^37^ The retroviral TR reporter was stably expressed in LNCaP as described to generate the LNCaP-TR cell line.^40^ LNCaP-TR-BoM was derived as detailed in Section 2.2 below. An *EPHA3*-ablated version of LNCaP-TR-BoM was derived as detailed in Section 2.9 below. LNCaP-RF, an androgen-refractory LNCaP subline was derived by extended propagation of LNCaP in androgen-deprived medium.^41^ All LNCaP-derived lines (including C4-2B), with the exception of LNCaP-RF, were cultured in RPMI-1640 medium (HyClone) with 10% fetal bovine serum (HyClone) and 1% Penicillin-Streptomycin (HyClone) supplemented. LNCaP-RF was cultured in similar conditions except with 10% charcoal stripped fetal bovine serum (Gentaur Genprice) supplemented instead of regular fetal bovine serum. DU-145 was cultured in DMEM with high glucose (HyClone) supplemented with 10% fetal bovine serum (HyClone) and 1% Penicillin-Streptomycin (HyClone). All lines were maintained at 37 °C in a humidified atmosphere containing 5% CO2. All cells were cultured for no more than twenty passages and tested monthly for mycoplasma-free status using a Mycoplasma Detection Kit (ABM, G238).

### 2.2. Derivation of LNCaP-TR-BoM

All animal procedures were approved by the Institutional Animal Care and Use Committee of the University of Notre Dame. LNCaP-TR cells were injected into the left cardiac ventricle of ten, 6-week-old male nude mice (Taconic, NCRNU-M) using established experimental metastasis methods.^40^ Lesion development was monitored longitudinally by bioluminescence imaging (IVIS Lumina) over an observation period of 190 days, using established luciferin dose and route.^40^ Only one animal developed a lesion in the middle spine, while the rest 9 mice did not develop detectable lesions. Visual inspection confirmed that the lesion was intraosseous rather than paraspinal soft tissue. The entire lesion was excised under sterile conditions, dissociated with 1 mg/ml collagenase IV (STEMCELL Technologies, 07427) for 1 h to prepare single cells. The recovered cells were propagated in culture and GFP^+^ cells were subsequently sorted with FACS to generate the LNCaP-TR-BoM cell line.

### 2.3. Two-dimensional culture and aggregate quantification

Cells were seeded at a density of 25000 cells per cm^2^ in standard 6-well tissue-culture plates (CELLSTAR) and imaged by brightfield microscopy on days 1, 3, and 5 after seeding (n = 3 (n refers to fields of view imaged within a given cell line’s well) per cell line per time point; 10x objective; scale bars 200 µm). The three fields of view imaged per well were determined by navigating approximately 1.75 cm from the center of the well in an upward, down-leftward, and down-rightward direction respectively. The same fields of view were followed longitudinally across the three time points. Cell clusters were defined as aggregates that visibly protrude from the monolayer and have a diameter of at least 50 µm. Cluster number per image and the projected area of each cluster were quantified. Imaging was done on an Etaluma Lumascope 720 (LS720). All image analysis and quantification were performed in ImageJ2 (Version: 2.16.0). For the 2D culture images, the Set Scale feature was first used to properly set the distances in the acquired images in the micrometer (µm) unit. Next, thresholding was performed for each image to clearly separate the cells from the background. Then, for each image, an ROI was drawn around each cell aggregate (clump) for the 2D culture images (the measuring tool in ImageJ2 was used to ensure that the diameter of each cell cluster quantified met the diameter cutoff of 50 µm). Finally, the Measure feature was used to obtain calculated numerical values for area (µm^2^) for each ROI (i.e. cell cluster). The number of cell aggregates (clumps) for the 2D culture images was equal to the number of ROIs drawn in each of those images.

### 2.4. Three-dimensional spheroid culture

For attachment-free culture, standard 96-well plates (CELLSTAR) were coated with poly(2-hydroxyethyl methacrylate) solution (0.6% poly-HEMA in 95% ethanol) to prevent adhesion.^42^ For the coating procedure, 50 µL of poly-HEMA solution was added to each well of the 96-well plate, and the plate was then left in a 37°C incubator for 3-4 days until all of the solution evaporated and then stored at room temperature. The poly-HEMA culture spheroids were imaged on days 2, 4, and 8 (n = 2 (n refers to replicate wells) per cell line per time point). The same wells were followed longitudinally across the three time points. For extracellular matrix-embedded culture, cells were seeded in Matrigel (Catalog #354230, Lot #03124002) and imaged on days 2, 4, and 6 (n = 3 (n refers to fields of view imaged within a given cell line’s well) per cell line per time point). The Matrigel stock solution (10.1 mg/mL) was used to carry out the Matrigel on-top 3D culture as described in Protocol 1 of Corning’s official Matrigel protocol (https://www.corning.com/catalog/cls/documents/application-notes/Application_Note_CLS-DL-AN-414_Matrigel_Matrix_3D_In_Vitro_Protocol.pdf). The same fields of view were followed longitudinally across the three time points. Imaging for both the poly-HEMA and Matrigel 3D cultures was done on a Leica DMi8 Inverted Microscope. All image quantification was performed in ImageJ2 (Version: 2.16.0). For the 3D culture images, the Set Scale feature was first used to properly set the distances in the acquired images in the micrometer (µm) unit. Next, thresholding was performed for each image to clearly separate the spheroids from the background. Then, for each image, an ROI was drawn around each spheroid for the 3D culture images. Finally, the Measure feature was used to obtain calculated numerical values for the area (µm^2^) and circularity (4π x area / perimeter^2^, where 1 denotes a perfect circle) for each ROI (i.e. spheroid). In the circularity analyses displayed in Figures 1G, 1J, and 5F, measurements from all time points were pooled within each cell line.

**Figure 1.**
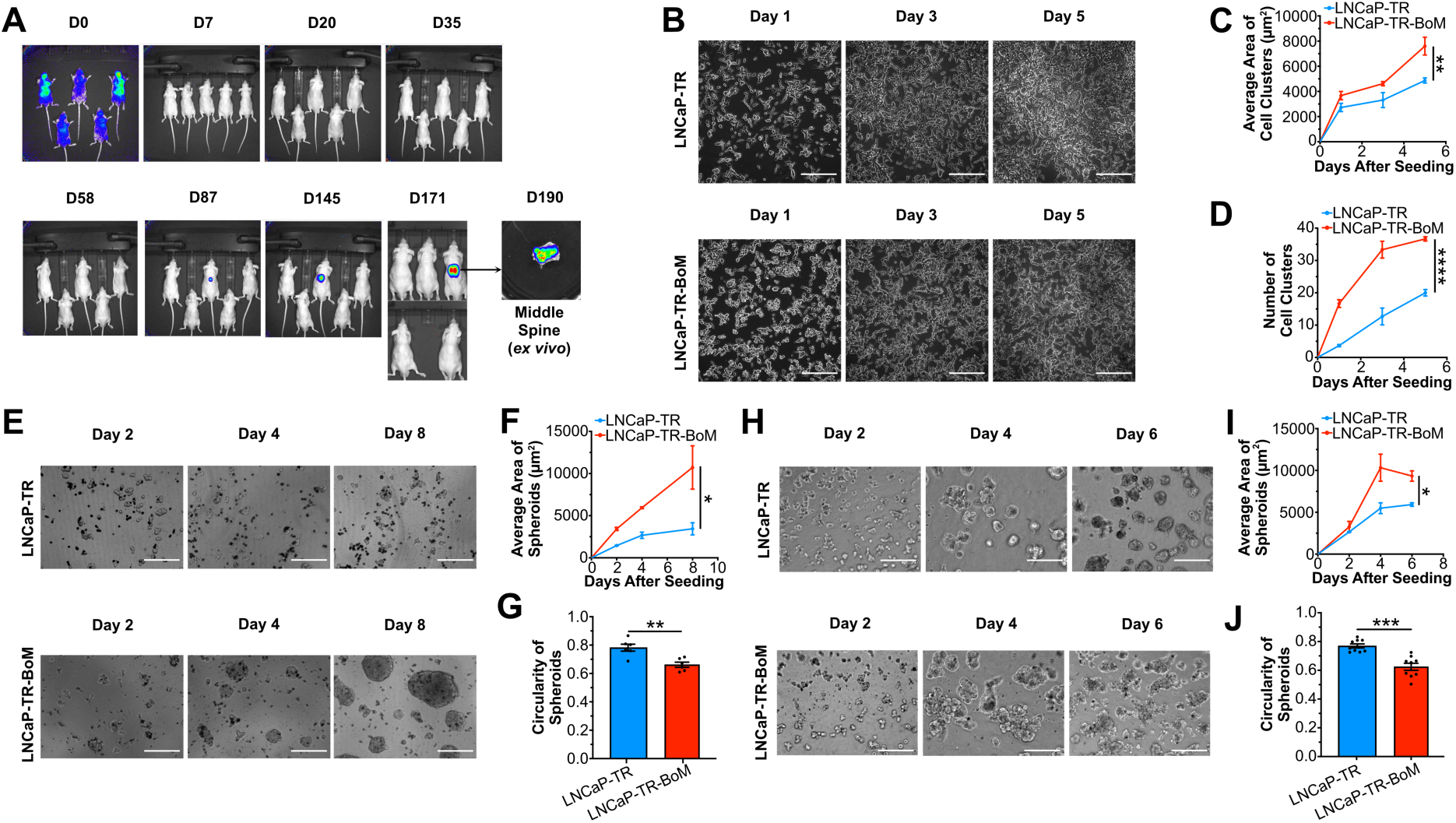
The *in vivo*-generated LNCaP-TR-BoM cell line demonstrates a more aggressive ability to form aggregates (clumps) and spheroids in 2D and 3D in vitro culture. (A) Bioluminescence imaging (BLI) images taken at the indicated time points over the span of 190 days after intracardiac inoculation of LNCaP-TR cells into 6-week-old male nude mice (n = 10 mice; only 5 mice (including the one mouse that developed a metastatic lesion) are shown), showing the middle-spine metastatic lesion from which the LNCaP-TR-BoM cell line was isolated and established. (B) Brightfield images of LNCaP-TR and LNCaP-TR-BoM cell lines in regular 2D culture at the indicated time points after seeding the cells (n = 3 per cell line per time point). (C) Quantification of average size (area) of cell aggregates (clumps) in each image from panel B. (D) Quantification of number of cell clusters in each image from panel B. (E) Brightfield images of spheroids formed by the LNCaP-TR and LNCaP-TR-BoM cell lines in 3D culture with poly-HEMA-coated 96-well plates at the indicated time points after seeding the cells (n = 2 per cell line per time point). (F) Quantification of average size (area) of spheroids in each image from panel E. (G) Quantification of circularity of spheroids in each image from panel E. (H) Brightfield images of spheroids formed by the LNCaP-TR and LNCaP-TR-BoM cell lines in Matrigel 3D culture at the indicated time points after seeding the cells (n = 3 per cell line per time point). (I, J) Same type of quantifications as in panels F and G but instead for each image from panel H. In (B), (E), and (H), all scale bars = 200 μm (objective lens magnification is 10x). In (C), (D), (F), and (I), data represent mean ± SEM; *p ≤ 0.05, **p ≤ 0.01, ***p ≤ 0.001, ****p ≤ 0.0001, two-way ANOVA with Bonferroni’s post hoc test with multiple comparison corrections. In (G) and (J), data represent mean ± SEM; *p ≤ 0.05, **p ≤ 0.01, ***p ≤ 0.001, ****p ≤ 0.0001, unpaired t-test with Welch’s correction. For (A), cell clusters were defined as cell aggregates (clumps) that visibly protrude up from the cell monolayer and that have a diameter of at least 50 µm. For circularity measures of the spheroids in (B) and (C), the scale is 0-1 where 1 means a perfect circle, and close to 0 means an elongated or highly irregular shape (images from all time points were pooled together for each cell line for circularity measures).

### 2.5. RNA sequencing and differential gene expression analysis

Total RNA (isolated with the Zymo Research R1054 Quick-RNA Miniprep Kit) was prepared from LNCaP-TR (n = 4), LNCaP-TR-BoM (n = 4), and LNCaP-RF (n = 3), where each biological replicate (n) was a separate cultured dish of a given cell line. RNA-seq was done with the HaploX company. Detailed quality control, library construction, other processing steps, and additional information (sequencing platform and instrument model, strandedness, reference genome build and annotation version, mapping and quantification of RNA-seq reads, etc.) can be found in the supplementary file (metadata.xIsx). Expression values in plots are reported as either counts per million (CPM), log2CPM, or z-score-normalized CPM. Prior to differential gene expression analysis, low-count genes (CPM < 1 in more than half of the compared samples) were filtered out. Normalization for library size and composition was done within the subsequent differential gene expression workflow. Differential gene expression analysis (carried out with the limma R package using the standard limma-voom workflow^43^) was assessed for LNCaP-TR-BoM relative to LNCaP-TR, and differentially expressed genes were determined using a log2FC (log2 of fold-change) absolute value cutoff of 1 and a Benjamini-Hochberg adjusted p-value cutoff of 0.05. When ranking by adjusted p-value, lower adjusted p-values correspond to higher rankings. The RNA-seq dataset used in this study can be accessed via NCBI Gene Expression Omnibus (GSE346590).

### 2.6. Quantitative reverse-transcription PCR

*KLK3*, *AR*, and *EPHA3* transcript levels were measured by quantitative reverse-transcription PCR (n = 3 per line for *KLK3* and *AR*; n = 6 per line for *EPHA3*). “n” here refers to technical replicates pipetted into separate wells of the 96-well qPCR plate. Reporting follows MIQE-consistent practice.^44^ After isolating total RNA using the EZ-10 Spin Column Total Mini-Preps Kit (Bio Basic Catalog #BS1361), All-In-One 5X RT MasterMix with gDNA Removal (abm Catalog #G592) was used to reverse transcribe the RNA into cDNA. We then performed qPCR using SYBR Green qPCR Master Mix (No ROX) (MedChemExpress Catalog # HY-K0523) on a CFX Connect Real-Time PCR Detection System (Bio-Rad). qPCR results were processed using the 2^ΔΔCT^ method to yield final values indicating relative RNA expression levels among different cell lines or treatment conditions. qPCR primer sequences are supplied in a supplementary Excel file. For each qPCR experiment, *GAPDH* or *RNA18SN5* was used as the housekeeping gene for normalization.

### 2.7. AR antagonist treatment

LNCaP-TR, LNCaP-TR-BoM, LNCaP-RF, DU-145, and C4-2B were treated for 3 days in 6-well plates with vehicle, 10 µM enzalutamide, or 10 µM darolutamide, after which *KLK3* transcript levels were measured as in Section 2.6 (n = 3 per cell line per condition). “n” refers to technical replicate wells on the qPCR plate when running the qPCR following AR antagonist treatment, RNA extraction, and reverse transcription. Enzalutamide (MedKoo Cat#: 201821) and darolutamide (MedKoo Cat#: 206514) were both obtained from MedKoo Biosciences, Inc. DMSO was used to reconstitute both drugs into 10 mM stock solutions. To reach the final working concentration of 10 µM, the stock solutions were diluted 1000x into the respective complete culture medium detailed in Section 2.1. The vehicle control therefore also contained a final DMSO % (v/v) of 0.1%.

### 2.8. Immunoblotting

Whole-cell lysates were resolved by SDS-PAGE and probed for AR, KLK3 (PSA), total EPHA3, phosphorylated EPHA3, and a loading control (β-actin in Figure 2C; GAPDH in Figure 4D and Figure 4E). Standard RIPA lysis buffer was used to harvest protein from cultured cells. After running SDS-PAGE to separate the proteins, the proteins were transferred to a nitrocellulose membrane, blocked with 5% milk (w/v), and then stained with primary antibodies and subsequently with secondary antibodies prior to chemiluminescence imaging with Clarity Max Western ECL Substrate (Bio-Rad #1705062) on a Bio-Rad ChemiDoc XRS+ imaging system. The primary antibodies used were Phospho-EphA3 Tyr779 (Cell Signaling Technology #8862), total EphA3 (Cell Signaling Technology #8793), PSA/KLK3 (Cell Signaling Technology #5365), Androgen Receptor (Cell Signaling Technology #5153), GAPDH (Proteintech #60004-1-Ig), and beta-Actin (Santa Cruz Biotechnology #sc-47778). The secondary antibodies used were Anti-rabbit IgG (HRP-linked) (Cell Signaling Technology #7074) and Anti-mouse IgG (HRP-linked) (Cell Signaling Technology #7076). Recommended dilutions to the final working concentration from the respective vendor’s websites were used for each antibody. More details about the primary and secondary antibodies used for immunoblotting are supplied in a supplementary Excel file.

**Figure 2.**
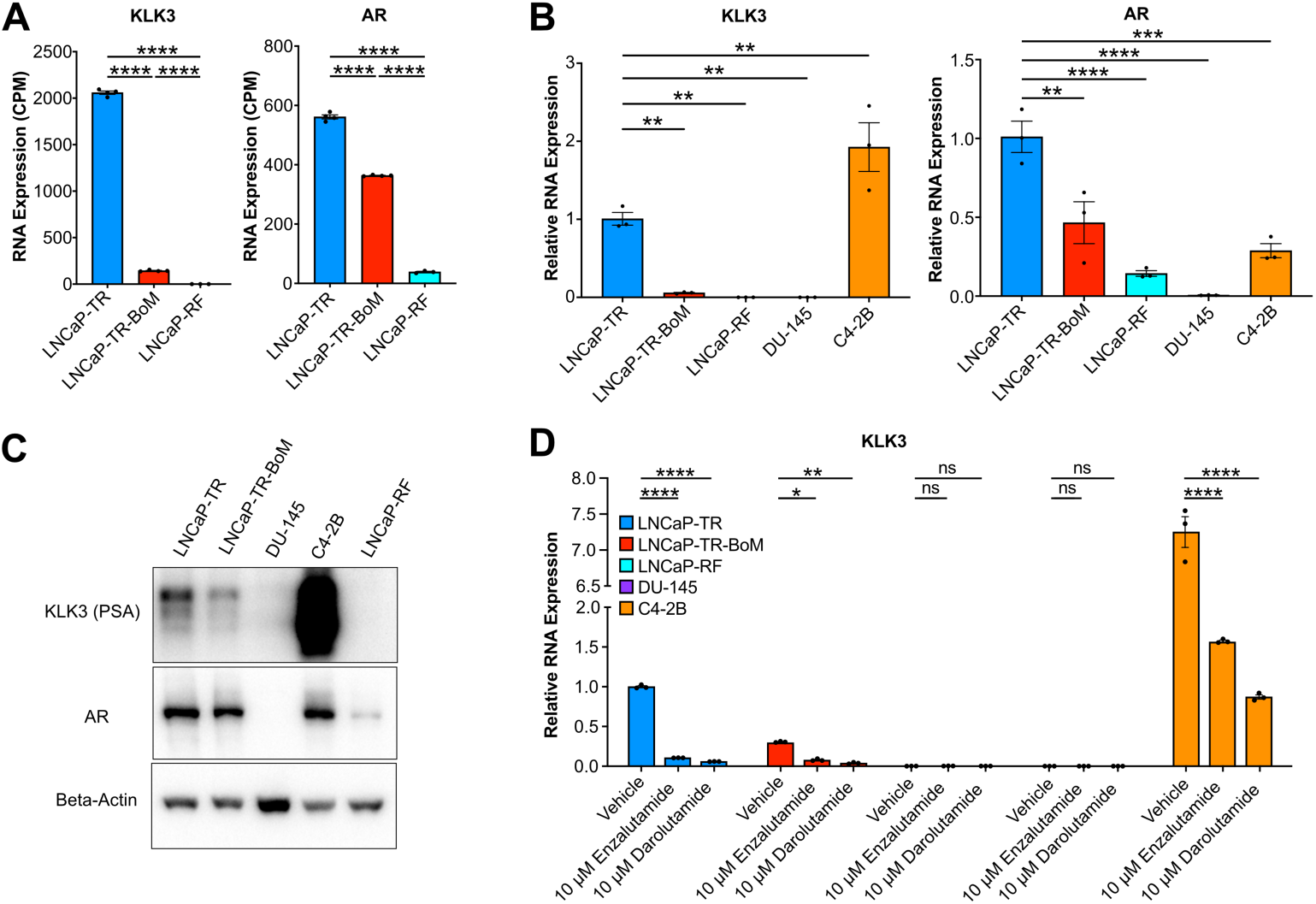
LNCaP-TR-BoM retains AR expression while AR pathway output is disproportionately reduced, indicating an intermediate AR-attenuated phenotype. **(A)** RNA expression levels (CPM-normalized) for *KLK3* and *AR* based on RNA-seq data for LNCaP-TR (n = 4), LNCaP-TR-BoM (n = 4), and LNCaP-RF (n = 3). (B) Relative RNA expression levels of *KLK3* and *AR* based on qRT-PCR for the indicated prostate cancer cell lines (n = 3 per cell line), normalized to LNCaP-TR. (C) Western blot to examine protein levels of KLK3 and AR in the indicated prostate cancer cell lines. (D) Relative RNA expression levels of *KLK3* (as determined by qRT-PCR) in the indicated prostate cancer cell lines after treatment with vehicle or AR antagonists at the indicated concentration for 3 days (n = 3 per cell line per treatment condition), normalized to the LNCaP-TR vehicle condition. In (A), data represent mean ± SEM; *p ≤ 0.05, **p ≤ 0.01, ***p ≤ 0.001, ****p ≤ 0.0001, one-way ANOVA with Tukey’s multiple comparisons test. In (B), data represent mean ± SEM; *p ≤ 0.05, **p ≤ 0.01, ***p ≤ 0.001, ****p ≤ 0.0001, one-way ANOVA with Dunnett’s multiple comparisons test. In (D) data represent mean ± SEM; *p ≤ 0.05, **p ≤ 0.01, ***p ≤ 0.001, ****p ≤ 0.0001, two-way ANOVA with Dunnett’s post hoc test with multiple comparison corrections. “ns” means not significant.

### 2.9. CRISPR-Cas9 editing of *EPHA3*

Three single-guide RNAs targeting *EPHA3* were evaluated in LNCaP-TR-BoM using CRISPR-Cas9.^45^ The three sgRNA sequences were GAACAGAGCAGCTGAGAAGG (#1), GTGGGAAGAGATCAGTGGTG (#2), and TTACCAGGTGTGCAATGTCA (#3). Delivery of these sgRNA sequences was done using lentiviruses generated using the Human EPHA3 sgRNA CRISPR/Cas9 All-in-One Lentivector set (Applied Biological Materials Inc. Catalog #19356111). Following selection with puromycin, the LNCaP-TR-BoM derivative generated with sgRNA #1, which lacked detectable EPHA3 protein by immunoblot, was designated as “*EPHA3* KO/Knockout” and used for all subsequent assays (heterogeneous transduced population was used rather than any particular colony-picked clonal population). LNCaP-TR-BoM cells that were not transduced with any *EPHA3*-targeting sgRNA were used as the matched control.

### 2.10. Public dataset analyses

Clinical primary prostate tumor and metastatic CRPC expression profiles were compared using the GSE35988 (GPL6480 platform) dataset from the Gene Expression Omnibus (GEO) via the PCaDB portal (http://bioinfo.jialab-ucr.org/PCaDB/).^46,47^ The Prostate Cancer Atlas^25^ was used to relate *KLK3* and *AR* RNA and protein levels to trajectory-inference pseudotime across clinical samples, and to compute pseudotime scores for the LNCaP-TR, LNCaP-TR-BoM, and LNCaP-RF transcriptomes (i.e. RNA-seq samples). Pseudotime scores for the LNCaP-TR, LNCaP-TR-BoM, and LNCaP-RF were subsequently normalized among these three cell lines’ RNA-seq samples (the lowest pseudotime score among the three cell lines’ RNA-seq samples was defined as 0 and highest pseudotime score defined as 1). *EPHA3* expression across established prostate cancer cell lines was examined in the Pan-cancer Cell Line Transcriptome Atlas (PCTA),^48^ in which each line is represented by many independently compiled RNA sequencing datasets; the n values reported for Figure S2B are numbers of compiled datasets, not biological replicates generated in this study. Pan-cancer *EPHA3* mRNA expression was examined in The Cancer Genome Atlas,^49^ and EPHA3 protein positivity across cancer types was examined in patient tissue microarray (immunohistochemistry/IHC) datasets from The Human Protein Atlas.^50,51^ Patients with “Medium” or “High” intensity IHC staining signal for EPHA3 in the “cancer_data.tsv.zip” data file (downloaded from The Human Protein Atlas) were considered to be positive for EPHA3 protein.

### 2.11. Pathway enrichment analysis

Over-representation analysis (ORA) and gene set enrichment analysis (GSEA)^52^ were applied to the differential gene expression analysis results from the “LNCaP-TR-BoM versus LNCaP-TR” pairwise comparison using clusterProfiler^53^ with Gene Ontology gene sets^54^ obtained from the Molecular Signatures Database.^55^ An adjusted p-value threshold of 0.25 was applied to the displayed pathway selection, and these results are treated throughout as exploratory and hypothesis-generating. The same pathway enrichment analyses were run for the differential gene expression analysis results from the GSE35988 dataset’s “mCRPC vs. Primary Tumor” pairwise comparison and overlapping enriched GO terms were identified. For both ORA and GSEA, the background gene set included all genes that were not filtered out due to having a low count (filtering threshold is described in Section 2.5). For GSEA, genes were ranked by log2FC.

### 2.12. Statistical analysis

Statistical tests are specified in each figure legend. Two-condition comparisons used unpaired t-tests with Welch’s correction; single-factor multi-condition (i.e. more than two conditions) comparisons used one-way ANOVA with Tukey’s or Dunnett’s multiple comparisons test; and two-factor multi-condition comparisons used two-way ANOVA with Bonferroni’s, Tukey’s, or Dunnett’s post hoc test. For Figure 3C and 3D, the multiple unpaired t-tests were conducted with Welch’s correction and with multiple comparison corrections using the Holm-Sidák method. Data are presented as mean ± SEM or as box-and-whisker plots as indicated. Significance thresholds are *p ≤ 0.05, **p ≤ 0.01, ***p ≤ 0.001, and ****p ≤ 0.0001. “ns” means not significant. All statistical analyses were performed using GraphPad Prism (Version 10.5.0).

**Figure 3.**
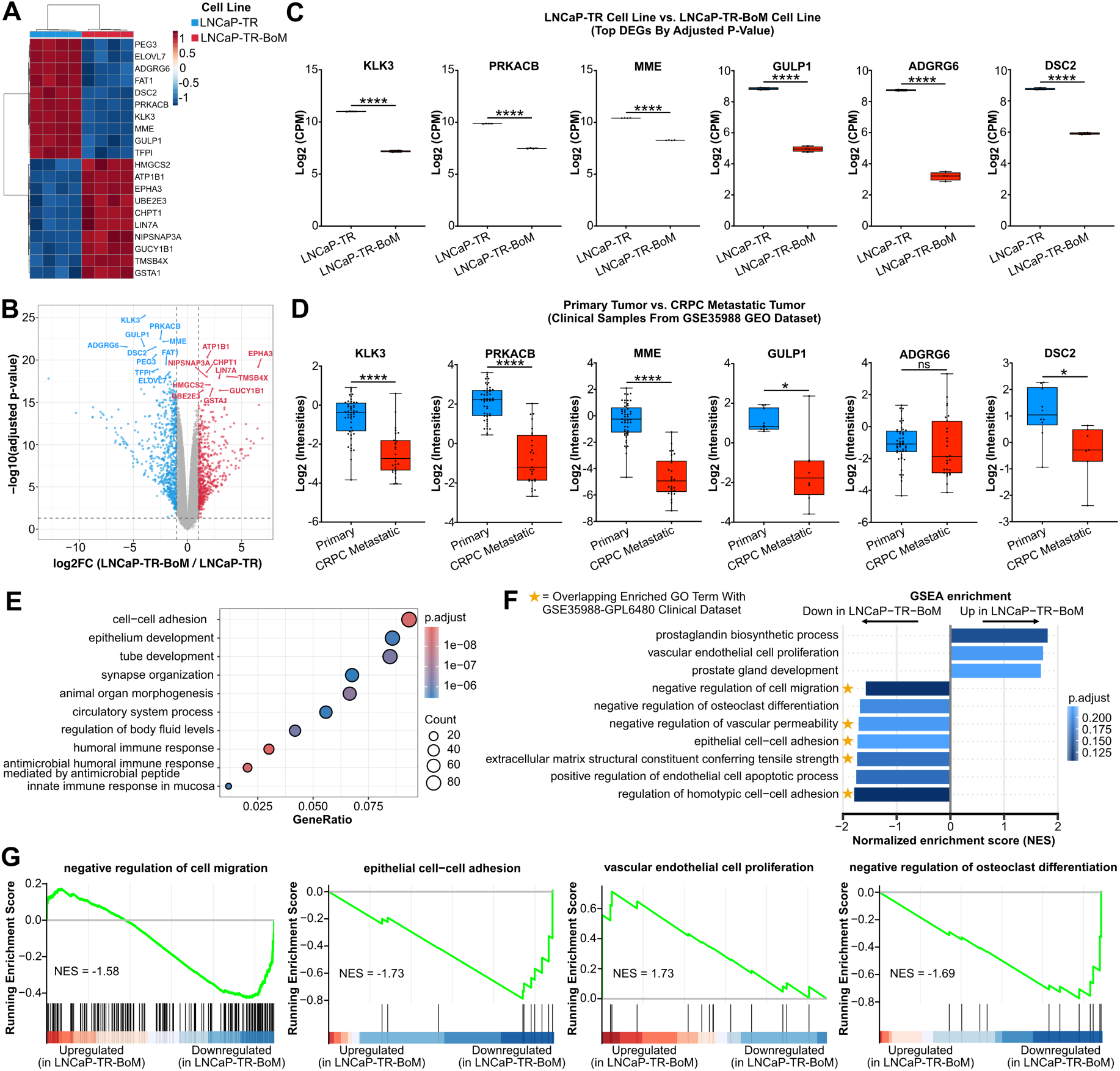
Bioinformatics analysis reveals a metastatically and clinically relevant transcriptomic signature for the LNCaP-TR-BoM cell line. **(A)** Z-score normalized gene expression clustered heatmap of the top 10 (by adjusted p-value) upregulated and downregulated differentially expressed genes (DEGs) in LNCaP-TR-BoM relative to LNCaP-TR. (B) Volcano plot of DEGs in LNCaP-TR-BoM relative to LNCaP-TR with the DEGs from panel A labeled; upregulated genes are colored in red, downregulated genes are colored in blue. (C) RNA expression levels based on RNA-seq data (log2-CPM-normalized) for the six top DEGs (ranked by adjusted p-value where lower adjusted p-value means higher rank) from the DGE analysis results for the LNCaP-TR vs. LNCaP-TR-BoM pairwise comparison; LNCaP-TR (n = 4) and LNCaP-TR-BoM (n = 4) cell lines were compared in the plots. (D) RNA expression levels based on microarray data (log2-intensity-normalized) from the GSE35988 GEO clinical dataset for the same six genes analyzed in panel C; primary tumor (n = 49 for *KLK3*, *PRKACB*, *MME*, and *ADGRG6*; n = 10 for *GULP1* and *DSC2*) and metastatic CRPC (n = 27 for *KLK3*, *PRKACB*, *MME*, and *ADGRG6*; n = 8 for *GULP1*; n = 7 for *DSC2*) clinical samples were compared in the plots. (E) Over-representation analysis (ORA) dot plot of enriched Gene Ontology (GO) terms for the LNCaP-TR-BoM vs. LNCaP-TR pairwise comparison; dot size denotes gene count and color denotes adjusted p-value. (F) Gene set enrichment analysis (GSEA) normalized enrichment scores (NES) for selected enriched GO terms for the LNCaP-TR-BoM vs. LNCaP-TR pairwise comparison; orange stars mark terms also enriched (in the same direction) in the GSE35988 clinical comparison (i.e. CRPC Metastatic vs. Primary); the sign (+ or -) of the NES indicates the enrichment direction of each GO term displayed. (G) GSEA running curve plots for selected enriched GO terms from (F). For pathway enrichment analysis results, an adjusted p-value cutoff of 0.25 was used. In (C) and (D), data are represented as box and whiskers plots; *p ≤ 0.05, **p ≤ 0.01, ***p ≤ 0.001, ****p ≤ 0.0001; in each panel, all six genes were compared via multiple unpaired t-tests with Welch’s correction and with multiple comparison corrections using the Holm-Sidák method. “ns” means not significant.

## 3. Results

### 3.1. *In vivo* derivation of LNCaP-TR-BoM and its altered multicellular organization in culture

To obtain a prostate cancer subline associated with skeletal colonization, we performed *in vivo* selection. LNCaP-TR cells were injected by the intracardiac route to 10 male nude mice and lesion development was tracked by bioluminescence imaging over approximately 190 days. Only one animal developed a lesion in the middle spine, and cells recovered from this lesion were established in culture as LNCaP-TR-BoM (**Figure 1A**). We first compared the growth behavior of LNCaP-TR-BoM and parental LNCaP-TR in standard monolayer culture (**Figure 1B**). Both lines remained adherent, but LNCaP-TR-BoM grew in a distinctly more aggregated manner. By day 5 after seeding, LNCaP-TR-BoM formed clusters with a mean projected area of approximately 7,600 µm^2^ compared with approximately 4,900 µm^2^ for the parental line, and formed approximately 37 clusters per field compared with approximately 20 for the parental line (**Figure 1C,D**).

Because enhanced cell-cell aggregation often predicts spheroid forming capacity, we next cultured both lines under attachment-free conditions in poly-HEMA-coated plates (**Figure 1E**). Over 8 days, LNCaP-TR-BoM formed spheroids that were larger on average and less circular than those of the parental LNCaP-TR (**Figure 1F,G**). The same directional differences were reproduced in Matrigel-embedded culture, which provides an extracellular matrix scaffold (**Figure 1H-J**). In three-dimensional culture systems, larger and less circular spheroid morphologies have been associated with poorly differentiated and invasive cell states, whereas smaller and more circular structures are typical of more differentiated cells.^56–59^ Therefore, these measurements are consistent with the hypothesis that LNCaP-TR-BoM represents a more aggressive phenotype compared with the parental cell line.

### 3.2. AR expression is largely retained in LNCaP-TR-BoM while AR pathway output is disproportionately reduced

We next characterized the AR axis, focusing on *AR* itself and on *KLK3*, which encodes prostate-specific antigen (PSA) and is a direct AR target gene widely used to gauge pathway output.^60^ RNA sequencing (RNA-seq) revealed an asymmetry between AR abundance and AR pathway output. Relative to parental LNCaP-TR, LNCaP-TR-BoM retained approximately 61% of *AR* mRNA but only approximately 7% of *KLK3* mRNA, an approximately nine-fold disparity between the two reductions; the androgen-refractory LNCaP subline LNCaP-RF showed very low *AR* and essentially undetectable *KLK3* (**Figure 2A**). Quantitative reverse-transcription PCR (qRT-PCR) across a broader panel reproduced this asymmetry: LNCaP-TR-BoM retained approximately 46% of parental *AR* but under 6% of parental *KLK3*, with both remaining clearly above the levels in LNCaP-RF and AR-negative cell line DU-145 (**Figure 2B**). Immunoblotting was qualitatively concordant, showing retained AR protein in LNCaP-TR-BoM together with a reduced PSA signal relative to the parental line, absent AR and PSA in DU-145, and a faint AR signal and undetectable PSA in LNCaP-RF (**Figure 2C**). We noted that the androgen-refractory LNCaP subline C4-2B expressed AR at mRNA and protein levels comparable with LNCaP-TR-BoM, but much higher PSA (**Figure 2B,C**). Therefore, the three LNCaP sublines (TR-BoM, RF, C4-2B) exhibit distinct AR-related expression features.

To test whether the residual AR pathway output in LNCaP-TR-BoM remains pharmacologically responsive, we treated the panel for 3 days with 10 µM enzalutamide or 10 µM darolutamide and measured *KLK3* (**Figure 2D**). Both antagonists suppressed *KLK3* in LNCaP-TR-BoM (approximately 78% with enzalutamide and approximately 87% with darolutamide, relative to its own vehicle control), in parental LNCaP-TR (approximately 90% and 94%), and in C4-2B (approximately 78% and 88%). *KLK3* remained at background in LNCaP-RF and DU-145, with no significant change. Because the fractional suppression achieved in LNCaP-TR-BoM was comparable to that in the parental line and in C4-2B, and because the assay endpoint is transcript abundance rather than cell growth, these data establish that AR pathway output in LNCaP-TR-BoM is attenuated but functional and druggable. They do not establish that LNCaP-TR-BoM is more resistant than the parental line to AR pathway-targeting treatment, a claim that would require growth or survival measurements under androgen deprivation and AR inhibition.

To place this phenotype in clinical context, we examined *KLK3* and *AR* along the prostate cancer progression trajectory defined by the Prostate Cancer Atlas, which orders clinical samples from normal tissue through primary tumors, metastatic AR-positive disease, and metastatic double-negative and neuroendocrine disease by trajectory-inference pseudotime.^25^ Both genes were lowest in the double-negative and neuroendocrine groups at the RNA and protein levels, consistent with loss of AR expression and output at the AR-null end of the trajectory, although the RNA relationship was not uniformly monotonic across the full pseudotime axis (**Figure S1A,B**). We then computed pseudotime scores from the transcriptomes of our three cell lines using the same portal. The three lines formed distinct transcriptomic clusters (**Figure S1C**), and their normalized pseudotime scores ordered LNCaP-TR-BoM between parental LNCaP-TR and LNCaP-RF (**Figure S1D**). We interpret this as relative transcriptomic positioning with respect to a clinical reference, not as evidence that LNCaP-TR-BoM is progressing over time toward a double-negative or neuroendocrine endpoint. Taken together, direct measurement and trajectory-based positioning indicate that LNCaP-TR-BoM occupies an intermediate, AR-attenuated state in which AR expression is largely preserved but canonical pathway output is substantially reduced.

### 3.3. Transcriptomic rewiring in LNCaP-TR-BoM overlaps with that of clinical metastatic disease

An intermediate AR-attenuated model is most useful if its molecular changes resemble clinical disease rather than an *in vitro* artifact. We therefore asked whether the transcriptomic differences between LNCaP-TR-BoM and parental LNCaP-TR are reflected in the differences between clinical mCRPC and primary prostate tumors in the Grasso et al (GSE35988) cohort.^46^

Differential expression analysis identified an extensive set of differentially expressed genes between the subline and its parent (**Figure 3A,B**). Six highly ranked downregulated genes, *KLK3*, *PRKACB*, *MME*, *GULP1*, *ADGRG6*, and *DSC2*, were all strongly and significantly reduced in LNCaP-TR-BoM (**Figure 3C**). In the clinical comparison, all six changed in the same direction, and five reached statistical significance (**Figure 3D**). We should note that this constitutes concordance for a selected set of top ranked (by adjusted p-value) genes. It does not establish transcriptome-wide equivalence between this cell line pair and clinical mCRPC, and we retain both non-significant comparisons for that reason. At the pathway level, over-representation analysis of the subline-versus-parent comparison highlighted Gene Ontology (GO) terms related to cell-cell adhesion, epithelium development, and tissue morphogenesis (**Figure 3E**). Exploratory GSEA identified changes in programs associated with cell migration, epithelial cell-cell adhesion, vascular endothelial biology, extracellular matrix structure, and osteoclast differentiation, several of which also appeared **(as indicated by annotations with orange stars in Figure 3F)** among the terms enriched in the clinical comparison (**Figure 3F,G**). We should note that these results are hypothesis-generating and are reported as candidate processes for future functional assessment rather than as established pathway-level equivalence. Furthermore, an important caveat applies to the framing of this comparison. LNCaP was itself established from a lymph node metastasis,^35^ so the LNCaP-TR and LNCaP-TR-BoM pair cannot be treated as a literal primary-to-metastatic progression series, and the parental line serves here as a non-skeletal, comparatively indolent reference rather than as a model of primary disease.

### 3.4. *EPHA3* is strongly upregulated in LNCaP-TR-BoM and is co-expressed with its ligand *EFNA5*

To test whether LNCaP-TR-BoM is usable as a discovery system, we selected a candidate from among its most significant differentially expressed genes. Among the ten genes with the lowest adjusted p-values, *EPHA3* showed the largest fold change and was strongly upregulated in the subline (**Figure S2A**). *EPHA3* encodes an ephrin type-A receptor tyrosine kinase; Eph receptors and their ephrin ligands regulate cell adhesion, migration, and tissue patterning and are frequently dysregulated in cancer, with context-dependent tumor promoting and tumor suppressing functions.^61^

Examination of the EPHA receptor and EFNA ligand families in our RNA-seq data showed that LNCaP-TR-BoM expressed, alongside high *EPHA3*, an appreciable level of *EFNA5*, a high-affinity EPHA3 ligand (**Figure 4A**). *EFNA5* was not itself upregulated in the subline relative to the parental line, but its presence in cells that also express high *EPHA3* raises the possibility of an autocrine or paracrine EPHA3 signaling interaction (not further tested in this study). Drastic *EPHA3* upregulation in LNCaP-TR-BoM indicated by RNA-seq was confirmed with qRT-PCR, showing an approximately 80-fold increase relative to parental cells (**Figure 4B,C**).

**Figure 4.**
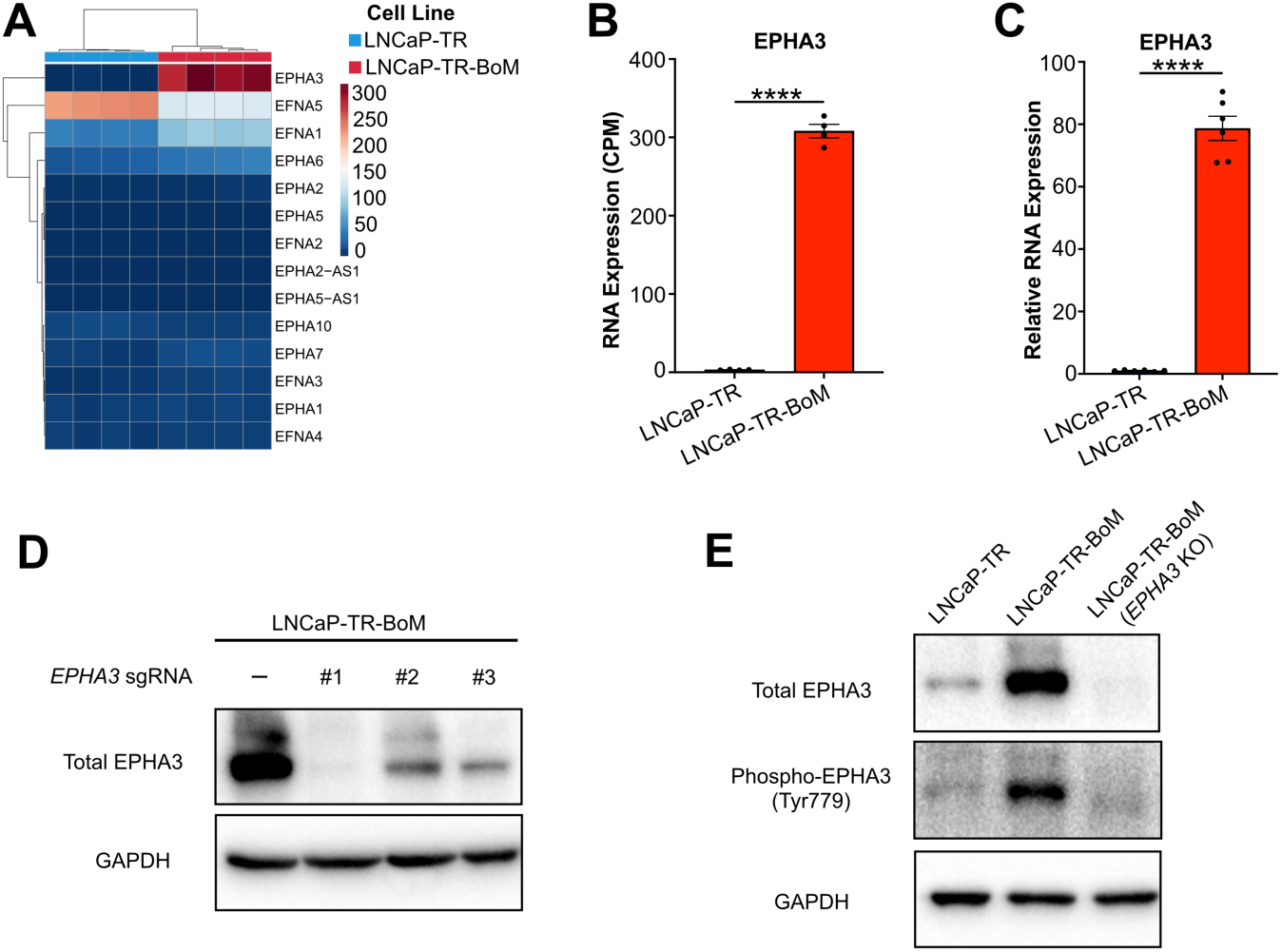
*EPHA3* is strongly upregulated by LNCaP-TR-BoM, prompting further investigation into its role via the generation of an *EPHA3*-knockout derivative of LNCaP-TR-BoM. (A) Heatmap showing the cpm-normalized expression levels of *EPHA* (ephrin receptors), *EFNA* (ephrin ligands), and associated antisense transcripts in the LNCaP-TR and LNCaP-TR-BoM cell lines based on RNA-seq samples. (B) RNA expression levels (CPM-normalized) for *EPHA3* based on RNA-seq data for LNCaP-TR (n = 4) and LNCaP-TR-BoM (n = 4). (C) Relative RNA expression levels of *EPHA3* in LNCaP-TR (n = 6) and LNCaP-TR-BoM (n = 6) based on qRT-PCR, normalized to LNCaP-TR. (D) Western blot of total EPHA3 protein in LNCaP-TR-BoM cells transduced with lentiviral vectors encoding Cas9 and sgRNA targeting *EPHA3* (three *EPHA3*-targeting sgRNA designs were tested (#1, #2, and #3); *EPHA3* sgRNA #1 was selected for use as “*EPHA3* KO” for subsequent data; “-” denotes the matched control where no *EPHA3*-targeting sgRNA was used). (E) Western blot of total EPHA3 protein and phosphorylated EPHA3 (Tyr779) protein in LNCaP-TR, LNCaP-TR-BoM, and LNCaP-TR-BoM (*EPHA3* KO) cells. In (B) and (C), data represent mean ± SEM; *p ≤ 0.05, **p ≤ 0.01, ***p ≤ 0.001, ****p ≤ 0.0001, unpaired t-test with Welch’s correction.

In the Pan-Cancer Cell Line Transcriptome Atlas (PCTA) compendium, *EPHA3* expression across established prostate cancer lines was highest in 22Rv1 and intermediate in C4-2B and C4-2, each above LNCaP, and was low in DU-145, PC3, and VCaP, with NCI-H660 comparable to LNCaP (**Figure S2B**). Elevated *EPHA3* is therefore a feature of some but not all castration-resistant lines. Cross-cancer analysis with TCGA and Human Protein Atlas data showed that prostate cancer was among the cancer types that exhibited the highest amounts of *EPHA3* expression at both the mRNA and protein levels (**Figure S2C,D**), supporting the clinical relevance of *EPHA3* in prostate cancer.

### 3.5. *EPHA3* disruption reduces aggregate and spheroid size and partially restores spheroid circularity

We generated three *EPHA3*-edited derivatives of LNCaP-TR-BoM using independent single-guide RNAs (sgRNAs). The derivative produced with sgRNA #1 lacked detectable EPHA3 protein, whereas residual protein remained after editing with sgRNA #2 and #3 (**Figure 4D**). Therefore, sgRNA #1 derivative was designated *EPHA3* KO and used for the functional assays. Immunoblotting confirmed that LNCaP-TR-BoM expressed substantially more total and phosphorylated EPHA3 than parental LNCaP-TR, and that neither signal was detectable in the *EPHA3* KO derivative (**Figure 4E**).

In monolayer culture, *EPHA3* KO reduced the mean projected area of cell clusters relative to LNCaP-TR-BoM, to a level statistically indistinguishable from parental LNCaP-TR (**Figure 5A,B**). Cluster number was numerically lower in the *EPHA3* KO derivative than in LNCaP-TR-BoM, but this difference did not reach significance (**Figure 5C**). In Matrigel culture, *EPHA3* KO spheroids had a smaller projected area than LNCaP-TR-BoM spheroids and were statistically indistinguishable from parental spheroids (**Figure 5D,E**). *EPHA3* loss in LNCaP-TR-BoM recovered spheroid circularity significantly yet did not reach the level of the parental cell line (**Figure 5F**). *EPHA3* disruption thus reverted spheroid size fully and spheroid shape partially toward the parental phenotype. These experiments identify EPHA3 as a contributor to the aggregation and spheroid phenotypes of LNCaP-TR-BoM and illustrate how this cell line can be used as a model system to nominate and test candidate genes that may contribute to the malignancy of AR-intermediate bone-metastatic prostate cancer.

**Figure 5.**
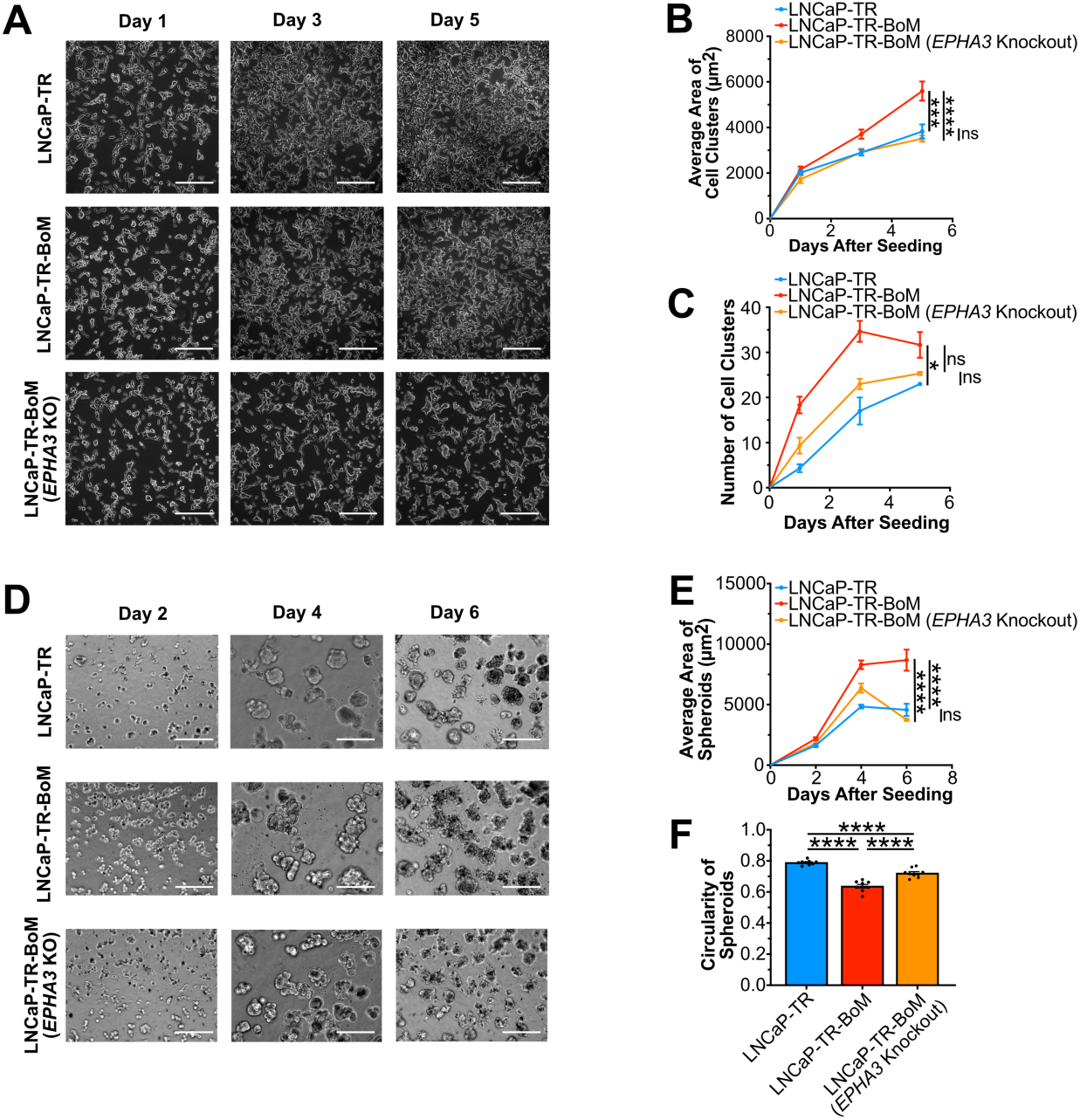
*EPHA3* disruption in LNCaP-TR-BoM lessens 2D aggregation, reduces 3D spheroid size, and partially restores spheroid circularity, providing support for the use of LNCaP-TR-BoM as a discovery system to test candidate genes. (A) Brightfield images of LNCaP-TR, LNCaP-TR-BoM, and LNCaP-TR-BoM (*EPHA3* KO) cell lines in regular 2D culture at the indicated time points after seeding the cells (n = 3 per cell line per time point). (B) Quantification of average size (area) of cell aggregates (clumps) in each image from panel A. (C) Quantification of number of cell clusters in each image from panel A. (D) Brightfield images of spheroids formed by the LNCaP-TR, LNCaP-TR-BoM, and LNCaP-TR-BoM (*EPHA3* KO) cell lines in Matrigel 3D culture at the indicated time points after seeding the cells (n = 3 per cell line per time point). (E) Quantification of average size (area) of spheroids in each image from panel D. (F) Quantification of circularity of spheroids in each image from panel D. In (A-F), scale bars, imaging parameters, and quantification methodology are the same as those described in Figure 1’s caption. In (B), (C), (E), data represent mean ± SEM; *p ≤ 0.05, **p ≤ 0.01, ***p ≤ 0.001, ****p ≤ 0.0001, two-way ANOVA with Tukey’s post hoc test with multiple comparison corrections. In (F), data represent mean ± SEM; *p ≤ 0.05, **p ≤ 0.01, ***p ≤ 0.001, ****p ≤ 0.0001, one-way ANOVA with Tukey’s multiple comparisons test. “ns” means not significant.

## 4. Discussion

We established LNCaP-TR-BoM, a prostate cancer cell line recovered from a spinal lesion that arose after intracardiac inoculation of TR-expressing LNCaP cells, and characterized it against its parental line. Three observations define the model. First, LNCaP-TR-BoM shows consistently altered multicellular organization: more numerous and larger aggregates in monolayer culture, and larger, less circular spheroids in two independent three-dimensional systems. Second, its AR axis is attenuated in a specific and informative way, with AR mRNA and protein largely retained but canonical pathway output, measured as *KLK3*, reduced by more than an order of magnitude, while remaining suppressible by AR antagonists. Third, its transcriptomic rewiring overlaps with that of clinical mCRPC relative to primary tumors for a set of highly ranked genes and for several candidate pathways relevant to bone metastasis.

The asymmetry between AR abundance and AR pathway output is, in our view, the most interesting feature of this model and the one that distinguishes it from the lines in common use. Widely used AR-high lines such as LNCaP, C4-2, C4-2B, 22Rv1, and VCaP sustain pathway output,^26–29^ while PC-3, DU-145, NCI-H660, and LASCPC-01 have neither AR nor its output.^30–32^ Clinical profiling of treatment-refractory mCRPC has established that AR expression and AR transcriptional activity can be uncoupled, and that intermediate and amphicrine states exist between the AR-high and AR-null extremes.^11,23^ LNCaP-TR-BoM appears to provide a tractable *in vitro* system in which that uncoupling is present in a defined, isogenic pair. Because the subline was never exposed to castration or to AR pathway-targeting drugs *in vivo*, the attenuated state arose through selection during skeletal colonization and *ex vivo* establishment rather than through direct pharmacological pressure, which distinguishes it from models generated by prolonged drug exposure.

Several limitations follow from the design of the study and should be stated explicitly. LNCaP-TR-BoM was derived from a single lesion in a single animal. Single-lesion derivation is shared by several classic prostate cancer lines, but reproducible bone tropism is a separate claim from cell line origin, and it requires re-inoculation of the subline and the parental line with quantitative comparison of skeletal colonization. The constitutive triple reporter makes such an experiment straightforward. Similarly, attenuated *KLK3* expression and pseudotime positioning are not measurements of growth dependence on AR. Whether LNCaP-TR-BoM is less dependent on AR for proliferation than its parent, and whether it is correspondingly less sensitive to androgen deprivation or AR antagonism at the level of cell growth, remains to be determined.

Our proof-of-concept experiments with *EPHA3* illustrate the intended use of the model while also showing its current limits. *EPHA3* was the largest-magnitude change among the most significant differentially expressed genes, was accompanied by expression of its high-affinity ligand *EFNA5*, and its disruption reduced aggregate and spheroid size and partially restored spheroid circularity. Ephrin signaling is bidirectional and strongly context-dependent. EPHA3 has been reported to contribute to prostate cancer growth and survival,^62^ to support tumor growth through stromal interactions in other settings,^63^ and to be functionally disrupted by somatic mutation elsewhere.^64^ In prostate cancer specifically, a YAP1-EPHA3 axis has been linked to cell contact, motility, cellular plasticity, and treatment response,^65,66^ EPHA-mediated signaling has been connected to stomatin regulation,^67^ and a circular RNA derived from the *EPHA3* locus has been reported to suppress progression,^68^ although a circular RNA is not evidence for an opposing function of the receptor protein itself. Our data place EPHA3 on the promoting side for aggregation and spheroid organization in this model. To reveal mechanism, the candidate EFNA5-EPHA3 loop is warranted to be formally tested for ligand dependence, with downstream effectors identified and the relevance of EPHA3 to bone colonization *in vivo* determined. We also note that, as a single cell line, LNCaP-TR-BoM cannot represent the inter-patient heterogeneity of clinical mCRPC, and its transcriptomic concordance with clinical samples, although encouraging, is correlative.

## 5. Conclusions

LNCaP-TR-BoM is a bone lesion-derived prostate cancer cell line that retains AR expression while showing markedly reduced AR pathway output, altered aggregate and spheroid organization, and transcriptomic changes that partly overlap with those of clinical mCRPC. Paired with its parental line, it offers a defined system for interrogating the intermediate portion of the AR phenotypic spectrum and for nominating and testing candidate genes, as illustrated here with *EPHA3*. Furthermore, the unique provenance and gene expression pattern supports the exploration of LNCaP-TR-BoM as a potential model of metastatic hormone-sensitive prostate cancer.

## Supporting information

metadata.xlsx

qRT-PCR Primers.xlsx

Western Blot (i.e. Immunoblotting) Antibodies.xlsx

## Author contributions

AC: Writing – review & editing, Writing – original draft, Visualization, Validation, Methodology, Investigation, Formal analysis, Data curation, Conceptualization. XL: Writing – review & editing, Supervision, Resources, Project administration, Investigation, Funding acquisition, Conceptualization.

## Acknowledgments

We want to thank the Lu lab members for their helpful suggestions during this work. We are grateful for the support from the core facilities at University of Notre Dame, including Freimann Life Science Center, Genomics and Bioinformatics Core Facility, and Integrated Imaging Facility. During the editing stage of this manuscript, the authors used Gemini in order to perform grammar checks and improve the linguistic clarity of the manuscript. The tool was not used to generate scientific content, interpret data, or draw conclusions. The authors take full responsibility for the content of the publication.

## Data availability

RNA-seq data of the cell lines are available at GEO with accession number GSE346590.

---

**Figure S1.**
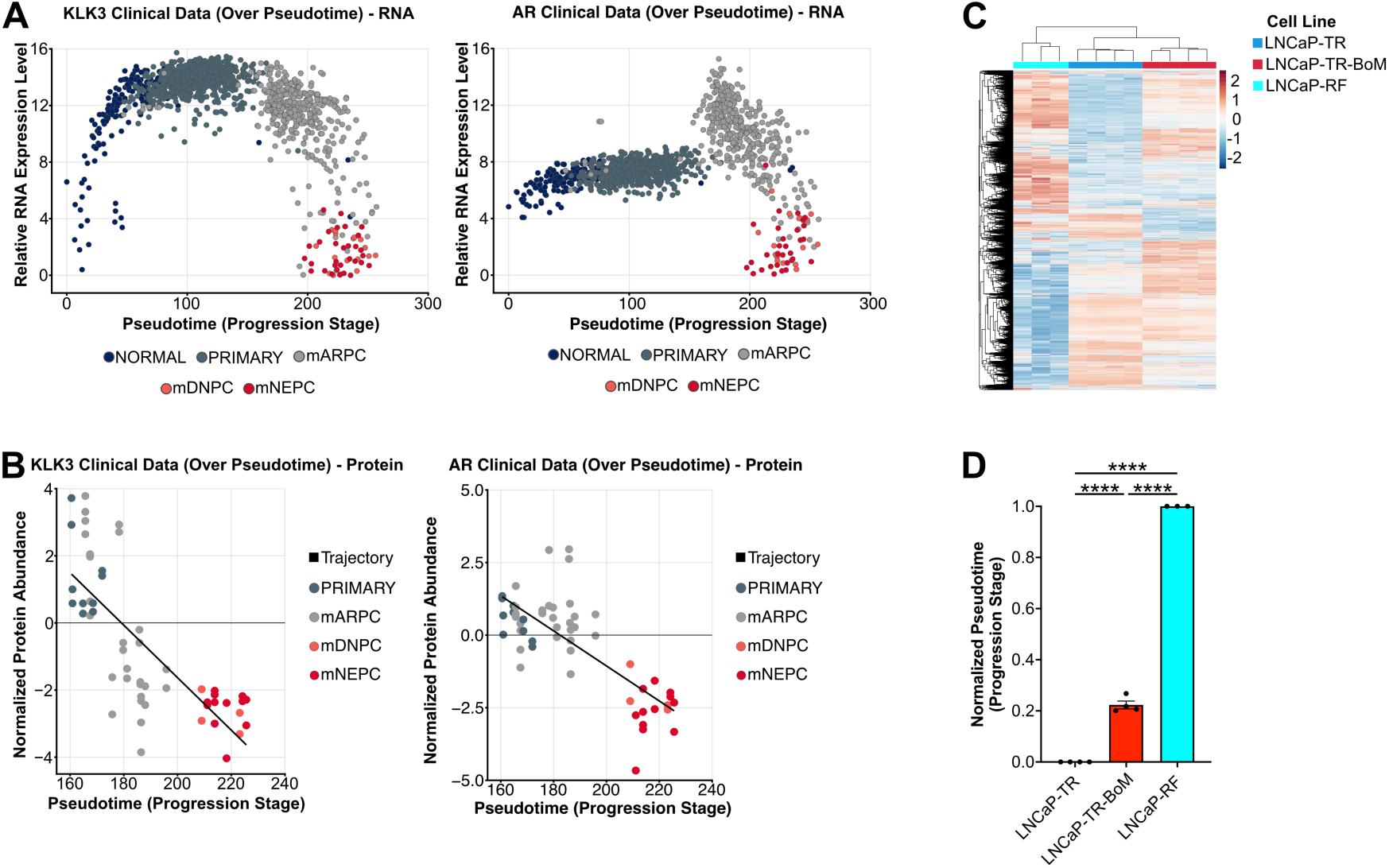
Positioning of the cell lines relative to a clinical prostate cancer transcriptomic trajectory. (A, B) Pseudotime cancer progression scatter plots depicting RNA (panel A) and protein (panel B) expression levels for KLK3 and AR in various clinical samples derived from prostate cancer patients (Prostate Cancer Atlas). (C) Z-score normalized gene expression clustered heatmap for the the LNCaP-TR, LNCaP-TR-BoM, and LNCaP-RF cell lines’ RNA-seq samples showing distinct transcriptomic signatures for the three cell lines. (D) Normalized pseudotime (progression stage) scores calculated from RNA-seq gene expression data (for LNCaP-TR, LNCaP-TR-BoM, and LNCaP-RF) indicating how advanced-stage each cell line’s overall phenotype is based on its transcriptomic signature. In (D), data represent mean ± SEM; *p ≤ 0.05, **p ≤ 0.01, ***p ≤ 0.001, ****p ≤ 0.0001, one-way ANOVA with Tukey’s multiple comparisons test.

**Figure S2.**
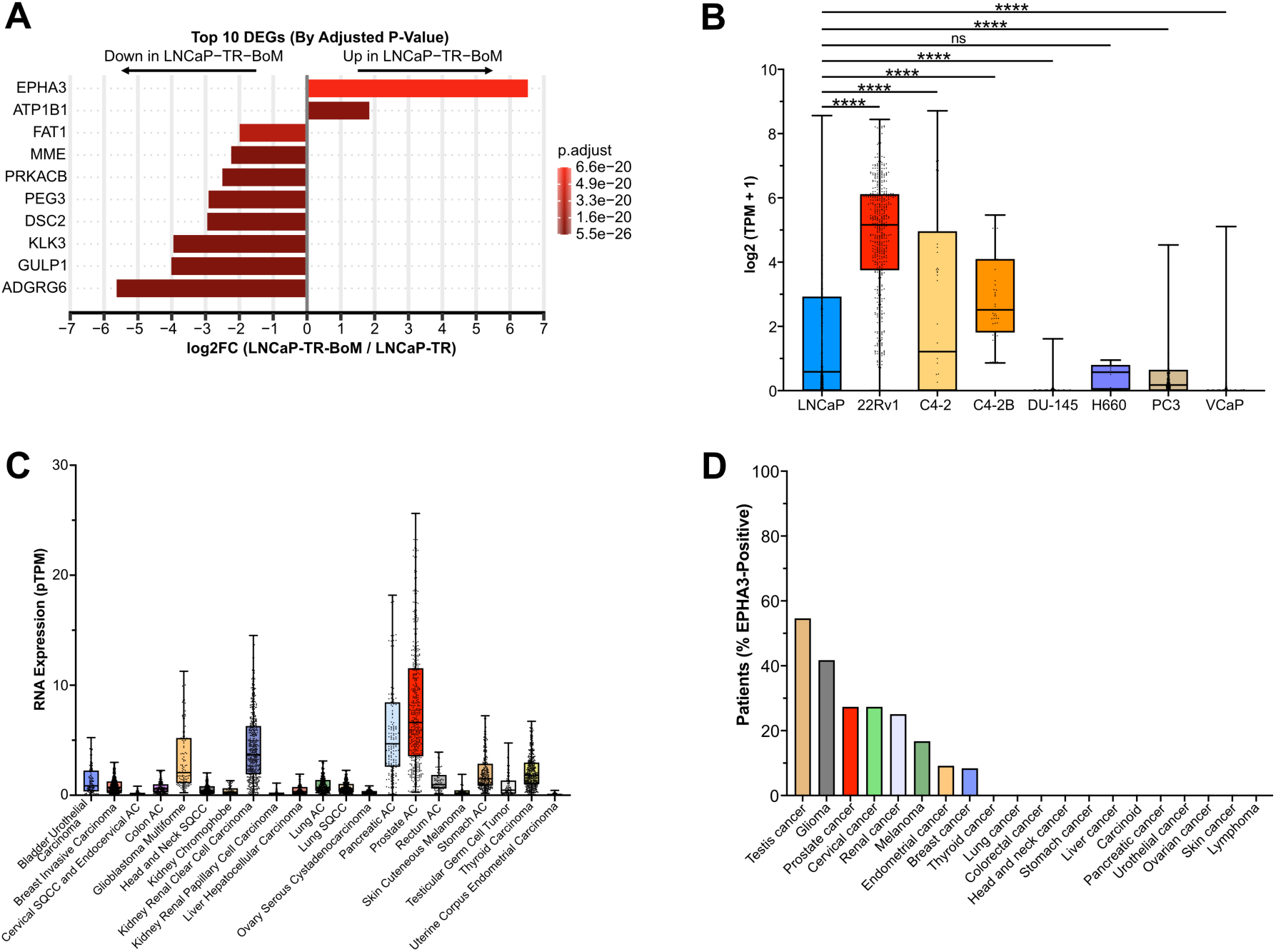
*EPHA3* ranking and clinical expression context. (A) Top 10 DEGs (by adjusted p-value) from the LNCaP-TR-BoM vs. LNCaP-TR pairwise comparison plotted with their log2FC values and adjusted p-values. (B) RNA expression levels (log2 (TPM + 1) normalized) of *EPHA3* in various prostate cancer cell lines based on compiled RNA-seq data from the Pan-cancer Cell Line Transcriptome Atlas (PCTA) dataset; LNCaP (n = 3330); 22Rv1 (n = 520); C4-2 (n = 51); C4-2B (n = 44); DU-145 (n = 716); H660 (n = 10); PC3 (n = 680); VCaP (n = 785). (C) RNA levels (pTPM-normalized) of *EPHA3* in TCGA clinical (patient-derived) samples across various types of cancer. (D) Percentage of clinical tissue samples positive for the EPHA3 protein across various cancer types based on patient tissue microarray (immunohistochemistry) datasets from The Human Protein Atlas. In (B), data are represented as box and whiskers plots; *p ≤ 0.05, **p ≤ 0.01, ***p ≤ 0.001, ****p ≤ 0.0001, one-way ANOVA with Dunnett’s multiple comparisons test.

